# Sex- versus apomictic-specific polymorphisms in the 5’UTR of APOLLO from *Boechera* shift gene expression from somatic to reproductive tissues in *Arabidopsis*

**DOI:** 10.1101/2023.08.29.555143

**Authors:** Maryam Honari, Joanne R. Ashnest, Timothy F. Sharbel

## Abstract

Among candidate genes underlying the control components of apomixis, APOLLO is known for its strong linkage to apomeiosis in the genus *Boechera*. The gene has “apo-alleles”, which are characterized by a set of linked apomixis-specific polymorphisms, and “sex-alleles”. All apomictic□*Boechera* genotypes are heterozygous for the apo/sex-alleles, while all sexual genotypes are homozygous for sex-alleles. In this study, native and synthetic APOLLO promoters were characterized by detecting the expression level of the β-glucuronidase (GUS) gene in *Arabidopsis*. Comparing various flower developmental stages in transgenic lines containing different constructs with 2 kb native transgenic lines revealed that changes to the APOLLO promoter causes shifts in tissue- and developmental-stage specificity of GUS expression. Importantly, several apomixis-specific polymorphisms in the 5’UTR change the timing and location of GUS activity from somatic to reproductive tissues. These synthetic data simulate a plausible evolutionary process whereby apomixis-specific gene activity can be achieved.

## Introduction

In contrast to sexual reproduction and segregation of traits in his peas, Mendel’s research on *Hieracium* illustrated a distinct kind of inheritance in which segregation was absent, and which was later identified as the parthenogenetic development of meiotically-unreduced egg cells (Bicknell et al, 2016; Mittelsten Scheid, 2022; van Dijk and Noel Ellis 2023). As with *Hieracium*, many plant species reproduce through apomixis, a natural form of asexual reproduction through which seeds are produced with embryos which are genetic clones of the mother plant (Hojsgaard and Hörandl, 2019; Underwood and Mercier, 2022).

While apomixis can arise via different mechanisms in natural plant populations (Hojsgaard and Hörandl, 2019; Underwood and Mercier, 2022), it fundamentally differs from sexual seed formation by the production of the meiotically-unreduced egg-cell (apomeiosis) which develops into an embryo without fertilization (parthenogenesis) in addition to endosperm development with or without fertilization of the central cell (Hojsgaard and Hörandl, 2019). Many apomicts are furthermore facultative (i.e., producing both sexually- and apomictically-derived embryos), and hence there is interplay between the reproductive pathways (Koltunow et al., 2011; Bicknell et al., 2003; Fehrer et al., 2007; Aliyu et al., 2010). Apomicts are often interspecific hybrids, which has led to the hypothesis that deregulation of genes underlying sexual seed formation underlies the genesis of apomixis (Carman, 1997). Furthermore, many apomictic plants and parthenogenetic animals are also polyploid, with implications for changes in gene expression and mutational load tolerance (among other traits) as being favorable for apomixis (Hojsgaard, 2018).

Apomixis has received much attention because of its potential ability to freeze complex hybrid genotypes (Tucker and Koltunow 2009). It does not exist in crop plants, and the introduction of apomixis from wild apomictic relatives into crops (Underwood et al., 2022), or via *de novo* induction would be highly disruptive to modern agriculture (Spillane et al., 2004; Underwood et al., 2022). Recent progress in inducing apomixis properties in lettuce and rice demonstrates the importance of research in this field to establish new tools to improve global food security (Underwood et al., 2022; Vernet et al., 2022).

The genus *Boechera* Á. Löve & D. Löve (Boecher’s rock cress; Brassicaceae)(Dobeš et al., 2006) is an ideal model system to study the evolution of apomixis, a major reason being that sexual and apomict plants can be compared genetically at the same ploidy level (Corral et al., 2013). The genus is characterized by diploid sexuals (basic chromosome number x = 7; 2n = 2x = 14), as well as diploid (2n = 2x = 14 or 15) and triploid (2n = 3x = 21) apomicts (Kantama et al., 2007; Alexander et al., 2013; Lovell et al., 2013). A highly reticulate pattern of genetic variation on inter- and intraspecific levels is reflective of numerous diploid and triploid apomictic lineages in geographically and genetically distinct populations (Rushworth et al., 2011; Rushworth et al., 2018; Kiefer et al., 2009), whereby rare haploid apomictic pollen can spread of apomixis factors to establishment new diploid apomictic lineages (Mau et al., 2021).

The introduction of apomixis-like reproduction in plants, using gene mutants (d’Erfurth et al., 2009), or through genetic constructs based upon candidate loci identified in different wild apomictic species (Underwood et al., 2022) demonstrate that multiple functional pathways are available to achieve asexuality. From a functional evolutionary perspective, deregulation of genes underlying sexual seed formation likely involves multiple loci which are simultaneously co-opted to induce apomixis, with changes to tissue-specificity and timing of gene expression leading to the switch from sexual to apomictic seed production in natural populations (Schmidt, 2020).

Here, we demonstrate that sex- vs. apomixis-specific polymorphisms in the 5’UTR of the apomixis candidate APOLLO gene (TGGCCCGTGAAGTTTATTCC)(Corral et al., 2013) can modify timing and location of gene expression in Arabidopsis. UTR-control of gene expression has been demonstrated to be significant in other plant species (Srivastava et al., 2018) as well as humans (Babu et al. 2021). For example, a novel motif in the 5’UTR of Big Root Biomass (BRB) gene modulates root biomass in sesame, whereby this motif in single and duplicated copies was responsible for high and low root biomass in two different accessions respectively (Dossa et al., 2021). Global spatial analysis of Arabidopsis natural variation revealed 5′UTR splicing of LATE ELONGATED HYPOCOTYL (LHY) in response to temperature (James et al., 2018). In addition, blue light exposure in Arabidopsis triggers variation in 5’UTR length resulting in differential transcription start site (TSSs) usage which in turn influences the post-transcriptional regulatory processes (Georgy, 2018; Yukio et al., 2018).

In *Boechera,* APOLLO is part of the DEDD 3′→5′ exonuclease superfamily (Corral et al., 2013), and the expression of its orthologue in *A. thaliana* (AT1G74390) has been reported in flowering stages, mature plant embryos, petal differentiation and expansion stages, plant embryo bilateral stages, plant embryo cotyledonary stages and plant embryo globular stages (https://www.arabidopsis.org). Sexual *Boechera* are homozygous for an APOLLO sex-allele, while apomicts have both apo- and sex-alleles which differ in sequence composition and structure (Corral et al., 2013). In *Boechera*, apo-alleles of APOLLO are highly expressed in developing apomictic ovules during the megaspore mother cell stage, while sex-alleles are not expressed in apomictic ovules (Corral et al., 2013). No APOLLO expression occurs in the ovules of sexual plants (Corral et al., 2013). Ovule-specific expression of APOLLO in apomictic *Boechera* is hypothesized to be associated with differences in the 5’UTR region which differs between apo- and sex-alleles, whereby the apo-allele 5’UTR is truncated and additionally contains an internal 20 bp insertion in comparison to the sex-allele (Corral et al., 2013). Here, we have employed a promoter-swap experiment between apo- and sex-allele specific sequences to investigate the influence of the apomixis-specific polymorphisms on the gene expression of this exonuclease in Arabidopsis.

## Materials and methods

### Plant materials and growth conditions

*Arabidopsis thaliana* (Columbia-0 ecotype) seeds were obtained from the Arabidopsis Biological Resource Centre (ABRC). Surface sterilised seeds were sown on MS plates containing ½ MS (Murashige & Skoog) medium (pH adjusted to 5.7 with 1M NaOH) with 0.8% phytoagar, stratified at 4°C for 48 h and then transferred to a phytochamber at 20°C with a light cycle of 16h light (white fluorescent bulbs, with a light intensity of 70 μmol m^−2^ s^−1^) and 8h dark. Two-week old seedlings were transferred to pots containing Sunshine mix 8 and grown in the same conditions.

### Construct generation and plant transformation

Native and promoter swap constructs were created using various combinations of sex- versus apo-specific polymorphisms identified in Corral et al. (2013; Figure 1, Supplemental Table 1). All primers used are described in Supplemental tables 2 and 3. All verified construct sequences have been submitted to Gene bank, as listed in Supplemental table 4.

**FIGURE 1.**
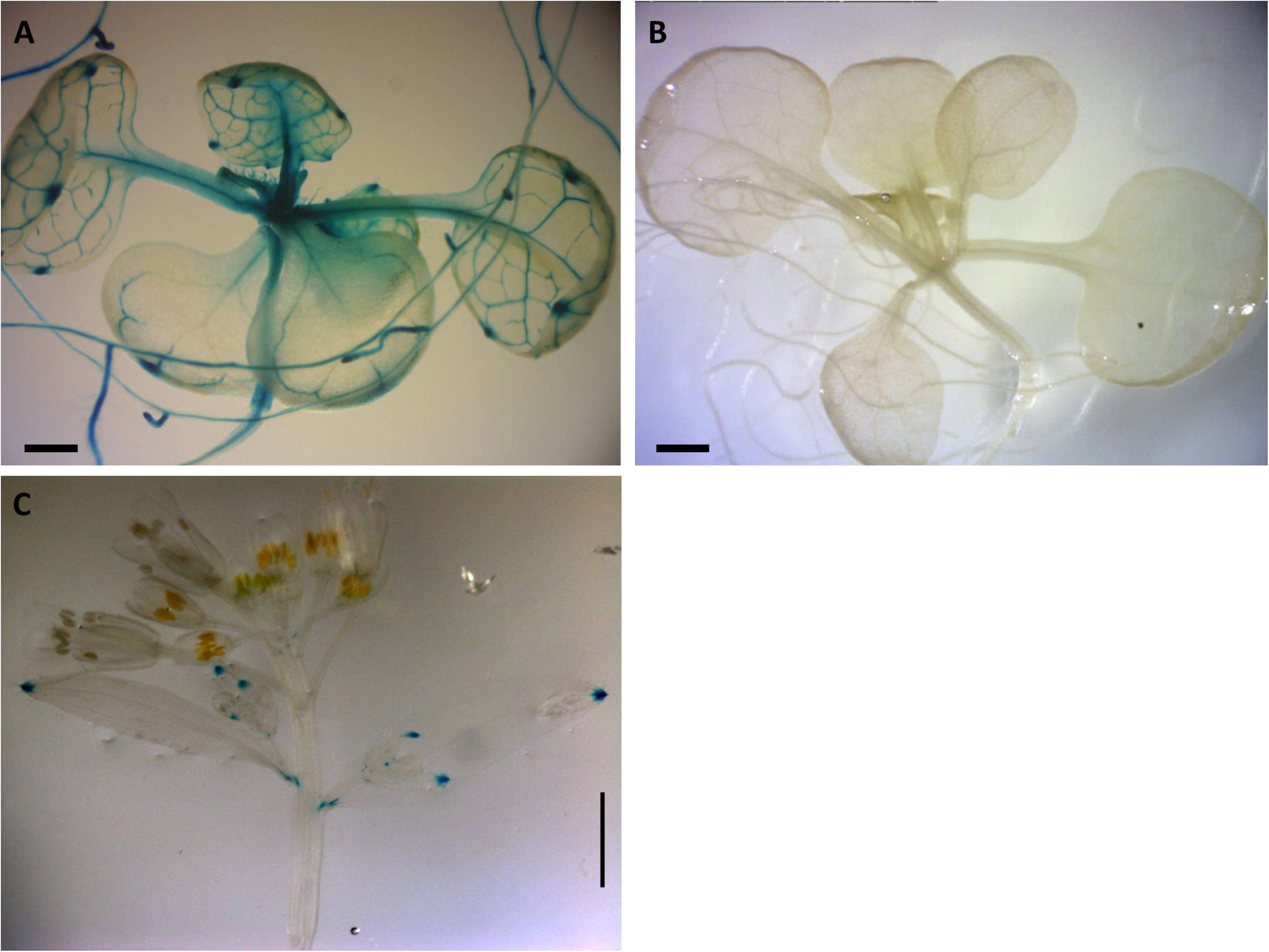
Construct pC1, pC2, pC3, pC4, pC5, pC6, and pC7. Constructs pC3 to pC7 created using SOE (Horton et al. 1989). For pC1, primer pairs of pAPOLLO AF-2 kb and pAPOLLO 1R were used to amplify 1,945 bp of DNA containing 71 bp of the 5’UTR of the Apo promoter containing the 20 bp Apo-specific insertion (yellow). For pC2, primer pairs of pAPOLLO SF -2 kb and pAPOLLO 1R were used to amplify 1,970 bp DNA containing 104 bp of the 5’UTR (gray). For pC3, primer pairs used to amplify 1,880 bp of the Apo promoter are shown in green, and primer pairs used to amplify 44 bp of the 5’UTR (gray) by deleting the 20 bp Apo-specific insertion (yellow) are shown in dark-blue. For pC4, primer pairs used to amplify 1,917 bp of the Sex promoter are shown in green, and primer pairs used to amplify 52 bp of Sex promoter by incorporating the 20 bp Apo-specific insertion (yellow) in dark-blue, with the 5’UTR of the Sex allele in gray. For pC5 primer pairs used to amplify 1,880 bp of the Apo promoter are shown in green, and primer pairs used to amplify 44 bp of the Apo promoter by incorporating a randomized 20 bp insertion (yellow) in dark-blue, with the 5’UTR of the Apo allele in gray. For pC6, primer pairs used to amplify 1,870 bp of the Apo-promoter are shown in green, and primer pairs used to amplify 104 bp of the Sex promoter in dark-blue, with the 5’ UTR of the Sex allele in gray, and the 20 bp Apo-specific insertion (yellow) deleted. For pC7, primer pairs used to amplify 1,865 bp of the Sex promoter are shown in green, and primer pairs used to amplify 71 bp 5’UTR of the Apo promoter carrying the Apo-specific insertion (yellow) is in dark-blue, with the 5’UTR of the Apo allele in gray color.

To build native *Boechera APOLLO* putative promoter constructs (C1 and C2), approximately 2 kb upstream of start codon, including the 5’UTR, was amplified from *B. stricta* (ES 718; sexual) and *B. divaricarpa* (ES 517; apomictic) genomic DNA using CloneAmp (Thermo Fischer Scientific) DNA polymerase. Specific primers were designed based on BAC apo- and sex-allele sequence libraries (Corral et al. 2013). Synthetic promoter splice variants (C3 to C7) were constructed using a modified “gene Splicing by Overlap Extension” method (SOE) (Horton et al., 1989). For C3, C4 and C5, modifications to the 5’UTR indel region were created by modifying the respective primer sequences (Supplemental figure 1). Construct pC5 was composed of the 2 kb native apo promoter plus the 5′UTR of the apo-allele, but the apomixis-specific 20 bp insertion sequence was randomized (but ratios of base pairs remained identical to the wild-type insertion; Figure 1). Resultant PCR products were purified and cloned into a pENTR™/D-TOPO™ entry vector via directional TOPO^®^ cloning, and incorporated into the pBGWFS7 GFP:GUS reporter vector (Nakagawa et al., 2007) using Gateway™ LR Clonase™. All constructs were verified using Sanger sequencing.

Binary vectors were transformed into the *Agrobacterium tumefaciens* strain GV3101 by electroporation, and eight-week old wild-type *Arabidopsis* Col-0 plants were transformed by floral dip (Clough and Bent, 1998). Transformed seeds were selected based on the method by Harrison et al. (2006). Two-week old glufosinate-resistant seedlings were then transferred to pots containing Sunshine mix 8, and presence of the transgene was confirmed by PCR of genomic DNA.

### Histochemical assay of GUS activity

Tissue-specific promoter activity for all constructs was assessed by β-glucuronidase (GUS) activity; seedlings and flowers from at least 10 different independent transformation events were examined per construct. GUS-staining was performed following the protocol of Jefferson (1987). Developmental staging was based on pistil and stigma maturity according to Smyth et al. (1990) and Christensen et al. (1997). We defined 5 developmental stages as in Rojeck et al. (2021) for analyses based upon pistil length, whereby the first stage (pre-meiotic stage) had a pistil length of 0.7 mm, and for the post-meiotic stages the pistil lengths were 1.0 mm, 1.5 mm, 2.0 mm and 3.0 mm. Among all of these stages the final stage is considered as post-fertilization stage (Rojeck et al., 2021)

Tissues were cleared following the protocol of Malamy and Benfey (1997). Samples were mounted in 50% glycerol on glass microscope slides. Among 10 different pre-assessed transgenic lines, 3 were used for microscopic analysis and photography, and *A. thaliana* var. Columbia that was not exposed to the clearing material was used as a control. Promoter activity in the leaves and flowers of *A. thaliana* was measured by a GUS histochemical assay at two- and five-days post-treatment in a minimum of three flowers per developmental stage (Smyth et al., 1990; Rojek et al., 2018). Images were obtained under a Zeiss Lumar.V12 Stereoscope and Olympus BX61 microscope and analyzed using Image J (https://rsb.info.nih.gov/ij/). To compare intensity of GUS positive areas, the linear region of interest (ROI) method was used, whereby a straight line is drawn through cell images to depict the main intensity of the color (Béziat et al., 2017). The original data was normalized by Min-Max method and analyzed using a two-way analysis of variance (two-way ANOVA). Finally, statistical differences were determined by Tukey test.

## Results

### (a) The 2kb native *Boechera* Apo promoter is expressed in the vascular tissues of *Arabidopsis*

Positive GUS activity for root, shoots, and leaves of two-week old seedlings for transgenic lines carrying the construct pC1 (2 kb native apo promoter) was observed in 33 lines of two-week old seedlings, each with 4 replicates (Figure 2). From this pattern (Figure 2a), three of the 33 lines were negative only for roots, shoots, or leaves, whereas 5 of 33 lines were negative only for roots (Supplemental figure 2). In contrast, no GUS activity was observed for 31 lines of two-week old seedlings transgenic for the construct pC2, carrying the 2 kb Sex native promoter (Figure 2b; Supplemental figure 2).

**FIGURE 2.**
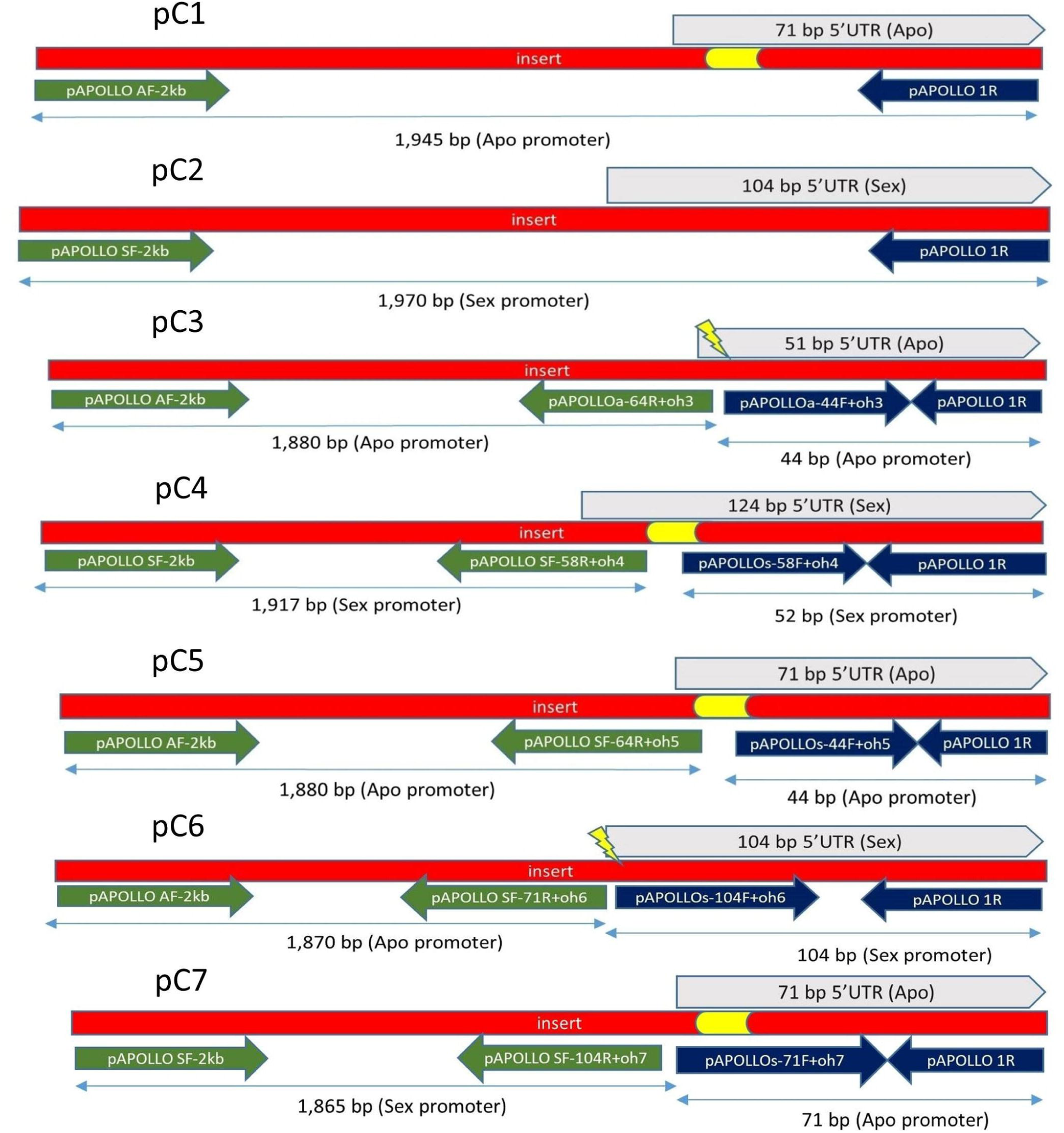
Expression patterns of Boechera native promoter GUS constructs in Arabidopsis. (A) GUS distribution of 2 kb native pApo promoter (pC1) in two-week old plant. (B) No GUS activity observed in 2 kb native pSex promoter (pC2). (A,B) Scale bar=0.5 mm (C) Positive GUS activity in leaf of six-week old Arabidopsis transgenic for pC1. Scale bar=2.0 mm (Zeiss Lumar.V12 Stereoscope).

There was positive GUS activity in leaves of six-week old *Arabidopsis* transgenic lines for pC1 (Figure 2c), and no GUS activity in any plants containing pC2 (Supplemental figure 2).

### (b) Removal of a 20 bp apo-specific insertion from the native 2kb apo-allele changes timing and tissue-specificity of expression

Construct pC3 (2kb native apo promoter plus 5’UTR with apo-specific insertion deleted; Figure 1) GUS activity was measured in 8 transgenic lines, over five different developmental stages (total of 40 flower buds). No GUS activity was observed in the pre-meiotic (0.7 mm) and post meiotic (1.0 mm) flower stages (Figure 3a, b, f, g), while positive GUS activity was observed for stages 3, 4, and 5 in the style and filament (including vascular tissue) with an increasing activity toward the final stages (Figure 3c, d, e, h, i, j; P≤0.05, Tukey test; Supplemental figure 3). Dispersed GUS activity in petals was excluded because they did not show a constant pattern for all samples. Thus, removal of the 20 bp apo-specific insertion from its native 2kb allele shifts expression from somatic to both male and female reproductive tissue.

**FIGURE 3.**
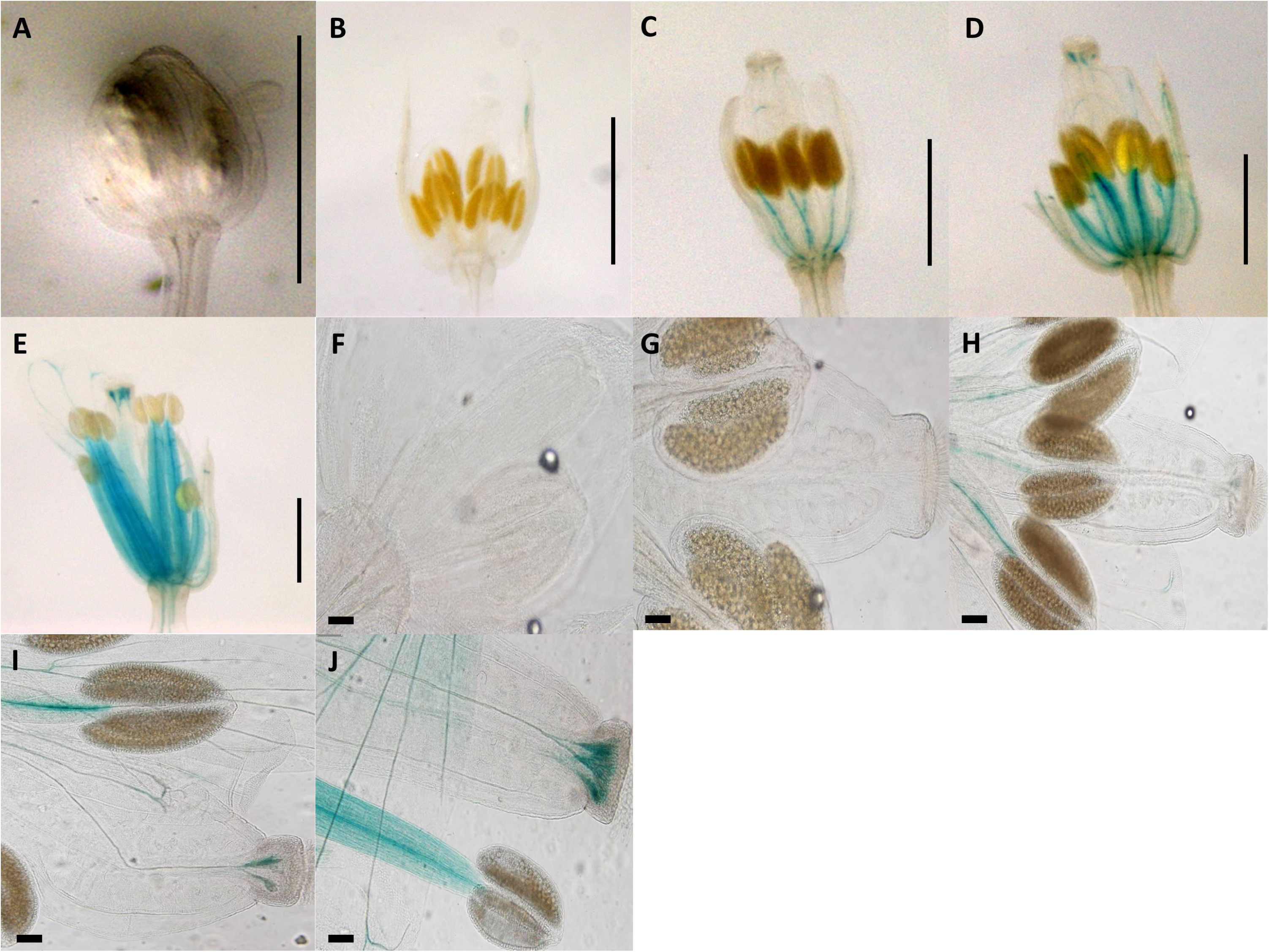
Expression patterns of synthetic promoter construct (pC3: GUS) in different developmental stages of Arabidopsis flowers. (A) 0.7 mm flower. (B) 1.0 mm flower. (C) 1.5 mm flower with GUS activity in filament. (D) 2.0 mm flower with GUS activity in style and filament. (E) 3.0 mm flower with GUS activity in style and filament. (A-E) Scale bar=1.0 mm (Zeiss Lumar.V12 Stereoscope), (F) 0.7 mm flower. (G) 1.0 mm flower. (H) 1.5 mm flower with GUS activity in style and filament. (I) 2.0 mm flower with GUS activity in style and filament. (J) 3.0 mm flower with GUS activity in style and filament, (F-J) Scale bar=100 μm (Olympus BX61 microscope).

### (c) A 20 bp apo-specific insertion does not induce expression in the 2kb native *Boechera* sex-allele

Construct pC4 carried the 2 kb native pSex promoter, the 5’UTR of the Sex-allele, with the 20 bp Apo-insertion added. The GUS activity for 7 transgenic pC4 lines, each of which had five different developmental stages (total of 35 flower buds) was measured, and no GUS activity was detected (Supplemental figure 4).

### (d) Randomizing the 20 bp apo-specific insertion but maintaining size induces anther and stigma expression of the native apo-promoter

Construct pC5 was composed of the 2 kb native pApo promoter plus the 5’UTR of the apo-allele, but a 20 bp randomized (i.e. same nucleotide composition but different sequence) insertion was substituted for the apomixis-specific 20 bp insertion (Figure 1). GUS activity was observed in the anthers and receptacle for the pre-meiotic stage, as well as some anther-specific GUS activity for stages 2 (Figure 4a, b, f, g; Supplemental figure 5). Style-specific GUS activity appeared in stages 3, and increased in intensity over stages 3, 4, and 5 (Figure 4c, d, e, h, i, j). GUS activity for the flower receptacle and petals were excluded for quantification because they did not show a constant pattern for all samples.

**FIGURE 4.**
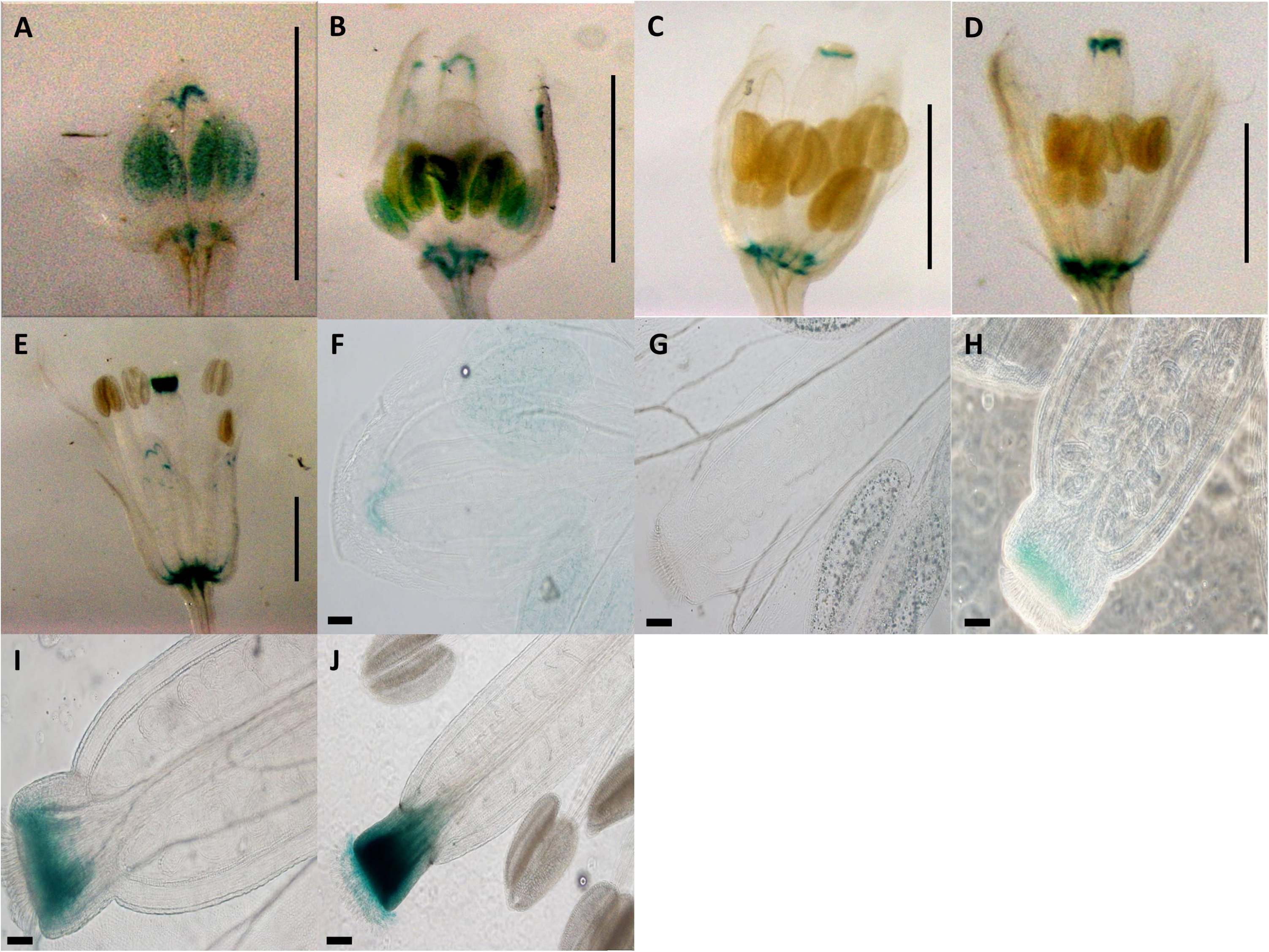
Expression patterns of spliced promoter construct (pC5: GUS) in different developmental stages of Arabidopsis flowers. (A) 0.7 mm flower with GUS activity in anther and receptacle. (B) 1.0 mm flower with GUS activity in anther and receptacle. (C) 1.5 mm flower with GUS activity in style and receptacle. (D) 2.0 mm flower with GUS activity in style and receptacle. (E) 3.0 mm flower with GUS activity in style and receptacle. (A-E) Scale bar=1.0 mm (Zeiss Lumar.V12 Stereoscope). (F) 0.7 mm flower with GUS activity in anthers. (G) 1.0 mm flower with GUS activity in anthers. (H) 1.5 mm flower with GUS activity in style. (I) 2.0 mm flower with GUS activity in style. (J) 3.0 mm flower with GUS activity in style. (G-J) Scale bar=50 µm

### (e) The apo-allele promoter with substituted sex-allele 5’UTR induces expression in polllen

Construct pC6 carried the 2 kb pApo promoter, with a swapped native 5’UTR of sex-allele (Figure 1), and was analyzed in 4 transgenic lines over five different developmental stages (total of 20 flower buds). Anther-specific GUS activity with varying intensity was observed for all developmental stages (Figure 5), with the strongest expression in stage 2 (Figure 5b; P≤0.05; Tukey test; Supplemental figure 6). We hypothesize that pollen activity is induced (Figure 5g to k), as is evidenced by post-dehiscence loss of activity (Figure 5d to e).

**FIGURE 5.**
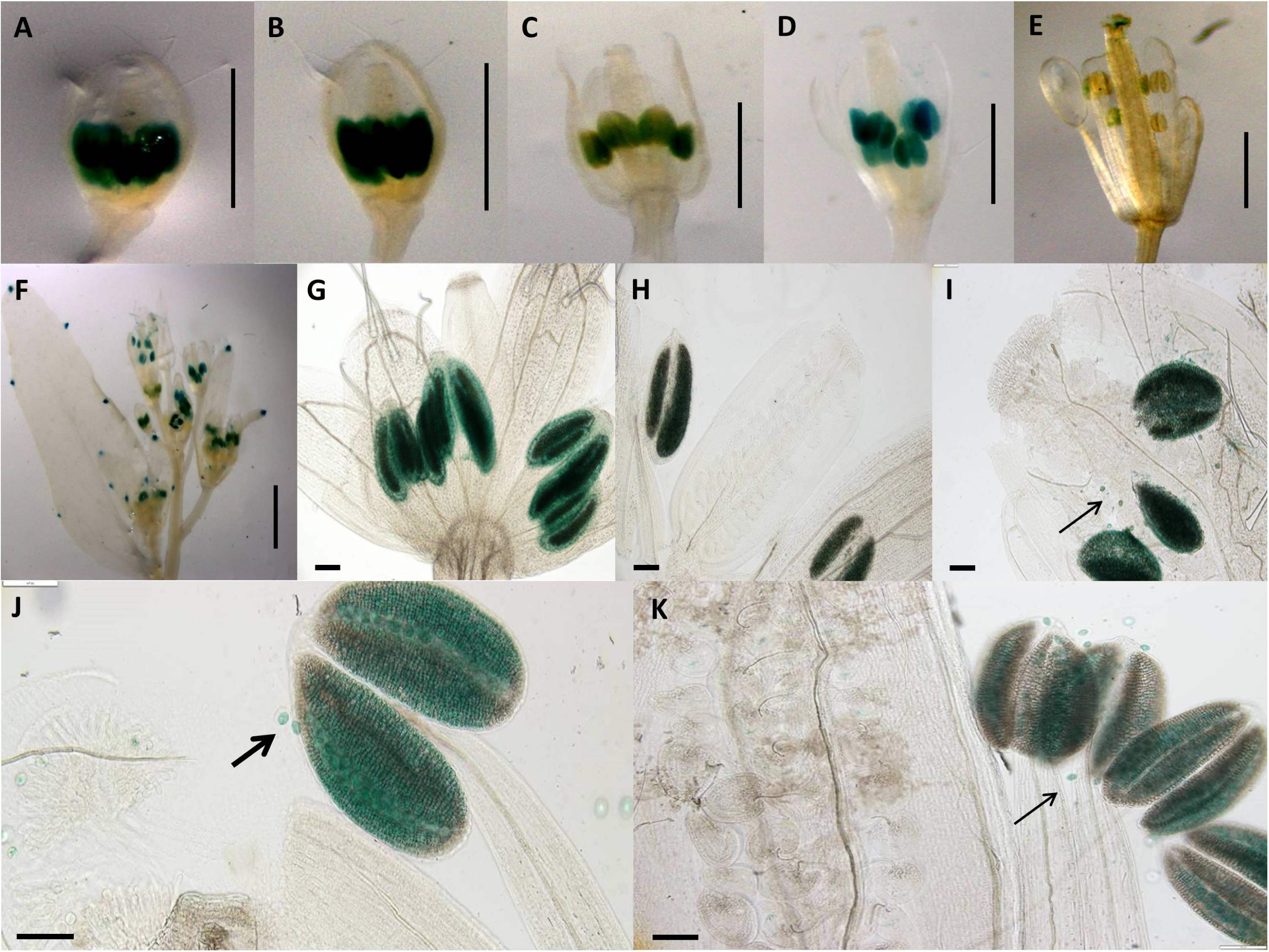
Expression patterns of spliced promoter construct (pC6: GUS) in different developmental stages of Arabidopsis flowers. (A) 0.7 mm flower with GUS activity in anthers, (B) 1.0 mm flower with GUS activity in anthers, (C) 1.7 mm flower with GUS activity in anthers (D) 2.0 mm flower with GUS activity in anthers, (E) 3.0 mm flower, (A-E) Scale bar=1.0 mm (Zeiss Lumar.V12 Stereoscope), (F) Arabidopsis inflorescence, scale bar=2.0 mm, (G) 0.7 mm flower with GUS activity in anthers, (H) 1.0 mm flower with GUS activity in anthers, (I) 1.5 mm flower with GUS activity in anthers, (J) 2.0 mm flower with GUS activity in anthers, (K) 3.0 mm flower, (G-K) Scale bar=50 µm (Olympus BX61 microscope). Pollen-specific GUS expression demonstrated by black arrows (I, J, K).

### (f) The sex-allele promoter with substituted apo-allele 5’UTR induces a burst of expression in both pollen and ovules in pre-zygote stages, followed by decreasing expression in zygotes

Construct pC7 carried the 2 kb sex promoter with a swapped apo-allele 5’UTR (Figure 1), and was analyzed in 6 transgenic lines over five developmental stages (total of 30 flower buds). GUS activity with a varied intensity was observed in the transmitting tract, ovules and anthers for the first two developmental stages (Figure 6a, b), while only ovule expression occurred in stage 3 (Figures 6c). Ovule expression began in stage one (Figure 6) and lasted for three stages (P≤0.05; Tukey test; Supplemental figure 7). In stage 4 (Figure 6d), expression in both tissues ceased.

**FIGURE 6.**
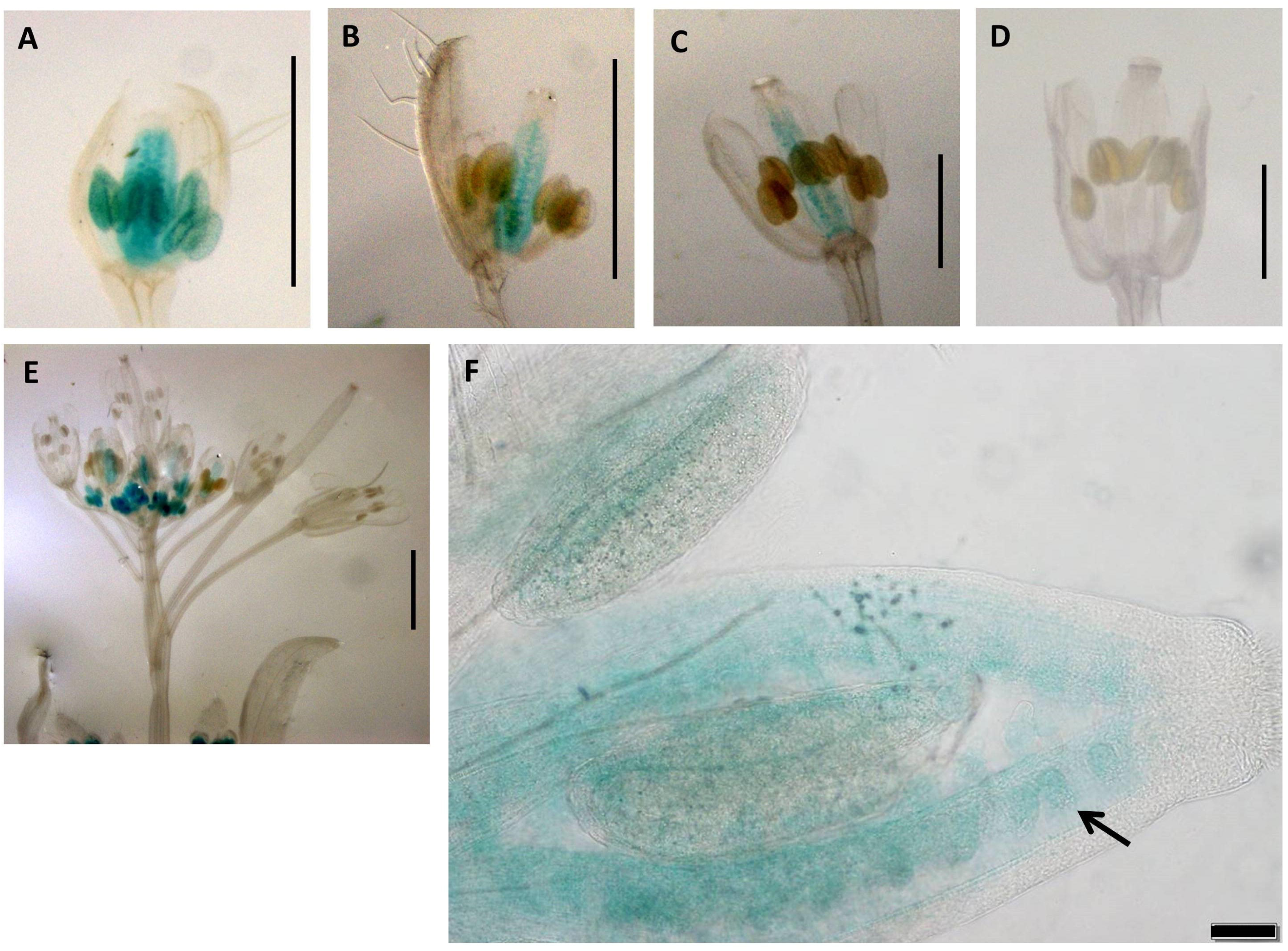
Expression patterns of promoter construct (pC7: GUS) in different developmental stages of Arabidopsis flowers. (A) 0.7 mm flower with GUS activity in transmitting tract, ovules and anthers, (B) 1.0 mm flower with GUS activity in transmitting tract and ovules, (C) 1.5 mm flower with GUS activity in and transmitting tract and ovules, (D) 2.0 mm flower, (A-D) Scale bar=1.0 mm (E) The Arabidopsis inflorescence, Scale bar=2.0 mm (Zeiss Lumar.V12 Stereoscope). (F) 1.0 mm flower showing expression in ovules (black arrows) Scale bar=50 μm (Olympus BX61 microscope).

## Discussion

A comparison of gene expression in the micro-dissected ovules of apomictic versus sexual *Boechera* led to the identification of the APOLLO gene. The heterozygous nature and presence of apomixis-specific polymorphisms were proposed to underlay gene expression differences between pre-meiotic ovules of apomictic versus sexual *Boechera* (Corral et al., 2013). Here, we investigated the nature of regulatory differences between native and synthetic APOLLO promoter GUS constructs in transformed Arabidopsis to understand functional differences between conserved sex-versus apo-allele polymorphisms in the 5′UTR region of the APOLLO gene. We have made no effort here to understand sequence variation in the 2 kb promoter regions of apo- and sex-alleles from multiple accessions of *Boechera*, as our focus was the 5′UTR region of the APOLLO gene. A comparison of the native promoter region sequences (constructs pC1 and pC2; Supplemental table 4) nonetheless demonstrates that these upstream regions also differ extensively, with potential implications for specific interactions in the promoter swap constructs which have not been tested here.

These data demonstrate that evolutionarily conserved apomixis-specific polymorphisms in the 5’UTR of the apo-allele in *Boechera* can cause changes to gene expression timing and location in *A. thaliana.* Importantly, coordinated expression in both male and female reproductive tissue can be achieved, thus providing a tool which can potentially be applied to reproductive genetics in Brassica plants.

### The native *Boechera* apo-allele promoter is active in the somatic tissues of *A. thaliana*

Arabidopsis transgenic lines for the 2 kb native Apo promoter demonstrate expression in the vascular tissues of roots stems and leaves, and expression ceases before the development of reproductive tissues (Figure 2). In contrast, no expression was observed for the 2 kb sex promoter in any Arabidopsis transformants.

In an analysis of the 5’UTR regions of APOLLO, Corral et al. (2013) showed that the *Boechera* 20 bp apomixis-specific insertion of the apo-allele shared a *SORLIP1AT* transcription factor binding site with the orthologous regions in *A. thaliana* and *Brassica rapa*, as well as a LIM1 transcription factor binding site with the orthologous regions in *A. thaliana*, while the sex-allele did not. Furthermore, the *Boechera* apo-allele transcripts are truncated at the 5’end of the 20 bp apo-specific insertion, while the sex-allele transcripts are approximately 35 bp longer (Corral et al., 2013). Finally, the apo-allele is characterized by exclusive expression in the ovules of apomictic *Boechera* (Corral et al., 2013), whereas in transformed sexual *A. thaliana* it is expressed in somatic and not reproductive tissues (Figure 2). In this case, we hypothesize that expression differences between the apo- and sex-alleles result from the different transcription factor landscape of the *A. thaliana* genome.

### The apomixis-specific insertion: length and sequence important for expression in male and female reproductive tissues

Compared to the native apo-allele promoter-5’UTR (Figures 1 and 2), removal of the 20 bp apomixis-specific insertion shifted expression from somatic to male and female reproductive tissues (Figure 3). pC3 showed GUS activity in filaments (including vascular tissue) and style in earlier and later stages of flower development respectively (Figure 3), and thus the absence of the 20 bp Apo-specific insertion changes the locality and timing of expression. Deletion of the 20 bp Apo-insertion thus likely leads to a change in the recognition site for regulating interacting elements that control expression in specific floral tissues.

Expression of the native *Boechera* apo-allele is not solely dependent on the apomixis-specific insertion, as insertion of the 20 bp apomixis-specific sequence into the native *Boechera* sex- allele produced no GUS activity in *A. thaliana* transformants (Supplemental figure 4). Alternatively, the 2 kb upstream promoter region of the sex-allele may not be long enough to drive expression.

Interestingly, the apomixis-specific insertion length is also important for expression in reproductive tissues, as randomization of the insertion sequence (with identical nucleotide composition) while maintaining length led to strong early expression in anther heads which stopped post fertilization. Post-fertilization was then characterized by growing strong expression in the stigma (Figure 4).

### Strong pollen-specific activity is induced by the apo-promoter/Sex-5’UTR construct

Transgenic pC6 lines carrying 2 kb of the apo promoter and the 5’UTR of the sex-allele (i.e., with no apomixis-specific 20 bp insertion) showed strong GUS staining in anthers in both pre- and post-meiotic floral buds (Figure 5). Closer examination of the anthers demonstrated that strong expression occurred in the pollen grains (Figure 5i, j). This expression ceased post-fertilization (Figure 5e).

### A coordinated window of activity in anthers and ovules during fertilization

Finally, transgenic pC7 lines carrying the 2 kb native sex promoter, 5’UTR of the apo-allele with the 20 bp apo-insertion (Figure 6; Supplemental figure 7) showed a window of GUS activity in both ovules and anthers in earlier stages of flower development which included the fertilization stage (Figure 6). As the native sex-allele promoter (pC2) showed no activity in Arabidopsis flowers (Figure 2), these data demonstrate that the apo 5’UTR can drive both male- and female-specific expression in *A. thaliana*.

Considering the orthologue of the APOLLO gene in *A. thaliana* (AT1G74390), the NAC45/86-DEPENDENT EXONUCLEASE-DOMAIN PROTEIN 3 (or NEN3), these data suggest that polymorphisms shared between the *Boechera* apo-5’UTR and that of *A. thaliana* are functionally involved with coordinating expression of this gene (or gene family) in male and female tissues during fertilization.

### The apomixis-specific insertion: size and sequence important for expression in male and female reproductive tissues

Removal of the 20 bp apomixis-specific insertion from the native apo-allele promoter-5’UTR (Figure 1) shifted expression from somatic (Figure 2) to male and female reproductive tissues (Figure 3). pC3 showed GUS activity in filaments and stigma in earlier and later stages of flower development respectively (Figure 3), and thus the absence of the 20 bp apo-specific insertion changes the locality and timing of expression. Deletion of the 20 bp apo-insertion thus likely leads to a change in the recognition site for regulating interacting elements that control APOLLO expression in specific floral tissues.

Expression of the native *Boechera* apo-allele is not solely dependent on the apomixis-specific insertion, as insertion of the 20 bp apomixis-specific sequence into the native *Boechera* sex-allele produced no GUS activity in *A. thaliana* transformants (Supplemental figure 4). Alternatively, the 2 kb upstream promoter region of the sex-allele may not be long enough to drive expression.

Interestingly, the apomixis-specific insertion length is also important for expression in reproductive tissues, as randomization of the insertion sequence while maintaining length led to strong early expression in anther heads which stopped post fertilization. Post-fertilization was then characterized by growing strong expression in the style (Figure 4).

### Strong pollen-specific activity is induced by the apo-promoter/sex-5’UTR construct

Transgenic pC6 lines carrying 2 kb of the Apo promoter and the 5’UTR of the sex-allele (i.e. with no apomixis-specific 20 bp insertion) showed strong GUS staining in anthers in both pre- and post-meiotic floral buds (Figure 5). Closer examination of the anthers demonstrated that strong expression occurred in the pollen grains (Figure 5i,j). This expression ceased post-fertilization (Figure 5e).

As the native apo-allele promoter drives expression in *Arabidopsis* while the native sex-allele promoter does not, the data so far demonstrate that the Apo-promoter upstream of the 5’UTR drives expression in *A. thaliana*, while the 5’UTR underlies male- or female-specific tissue expression.

### The 20 bp apo-specific insertion may bind a repressor in meiotic (sexual) cells

The similarities between the 5′UTR sequences of the *Boechera* apo-allele, *A. thaliana* and *B. rapa* suggest that the apo-allele is ancestral compared to the sex-allele, and that deregulation of APOLLO in apomicts likely underlies its expression in pre-meiotic tissues (Corral et al. 2013). This is supported by the data presented here, whereby in *A. thaliana*, a sexually reproducing species, removal or scrambling of the apo-specific 5′UTR insertion and neighboring sequence induced expression in both male and female reproductive tissue. Together, these data suggest that the apo-5′UTR encodes a sequence-specific binding site for a transcriptional or translational repressor, one that may be specifically expressed in sexual meiotic cells.

### Hybridization and differential gene expression between apomictic versus sexual *Boechera*

The elevated heterozygosity in diploid apomictic *Boechera* can be explained by widespread hybridization associated with the transition from sexuality to asexuality, or it could be a byproduct of apomixis itself. In apomictic individuals the overall heterozygosity is expected to increase over generations due to the lack of recombination and mutation accumulation (Meselson effect; Hojsgaard and Hörandl, 2015). Clear evidence for the hybrid origin of highly heterozygous apomicts has been shown (Beck et al., 2012), in addition to mutation accumulation on a genomic level in apomictic *Boechera* in conserved non-coding sequences (Lovell et al., 2017). Finally, rare haploid pollen produced by apomictic *Boechera* can transfer the complete apomictic trait in a single generation in both intra- and interspecific crosses onto emasculated female plants (Mau et al., 2021). Taken together, these data demonstrate that the conserved apo-allele of *Boechera* is found with different sexual alleles on a broad geographic scale, not to mention different genomic “contexts” considering the nuclear and cytoplasmic genomes in independently derived apomictic lineages (Kiefer and Koch, 2012; Kiefer et al., 2009).

Here we show that conserved polymorphisms in the native *Boechera* apo-allele 5’UTR are functionally associated with driving gene expression in both male and female tissues during important reproductive stages (e.g. gametogenesis and fertilization). The interspecific “collision” of different sexual genomes may produce apomictic *Boechera* with structural changes such as homeologous chromosome substitutions and aneuploidy (Kantama et al., 2007). Extending divergence to regulatory genes, apomixis could result from the novel pattern of gene expressions created by the interaction of divergent transcriptional regulators (Carman, 2001), and is supported by the data reported in this study.

## Supporting information

C:\Users\Administrator\Desktop\Revised paper for pre-print\Supplemental data.doc

## Acknowledgements

Experimental work was funded by a grant from the Global Institute of Food Security at the University of Saskatchewan to Timothy F. Sharbel. The authors thank Marco Pellino, Andres Posso-Terranova, Zahida Irin, Nazmul Hasan, Angie Li, Pierre-Luc Pradier for help with greenhouse operations and plant care, as well as laboratory assistance.

