## Supplementary material for "Sex- versus apomictic-specific polymorphisms in the 5’UTR of APOLLO from *Boechera* shift gene expression from somatic to reproductive tissues in *Arabidopsis*": C:\Users\Administrator\Desktop\Revised paper for pre-print\Supplemental data.doc

**Protocol for SOE**

These five different spliced promoters were prepared using a modified protocol based upon the “gene Splicing by Overlap Extension” method (SOE) (Horton et al*.*, 1989). This method allows segments from two different genes to be recombined or “spliced” together by overlap extension in which the copy of a DNA strand developed by the first PCR reaction can then serve as a template for an extension from a second primer in the opposite orientation (Supplemental figure 1). This method enabled the production of different constructs which contained different components of the apo- or the sex-allele.

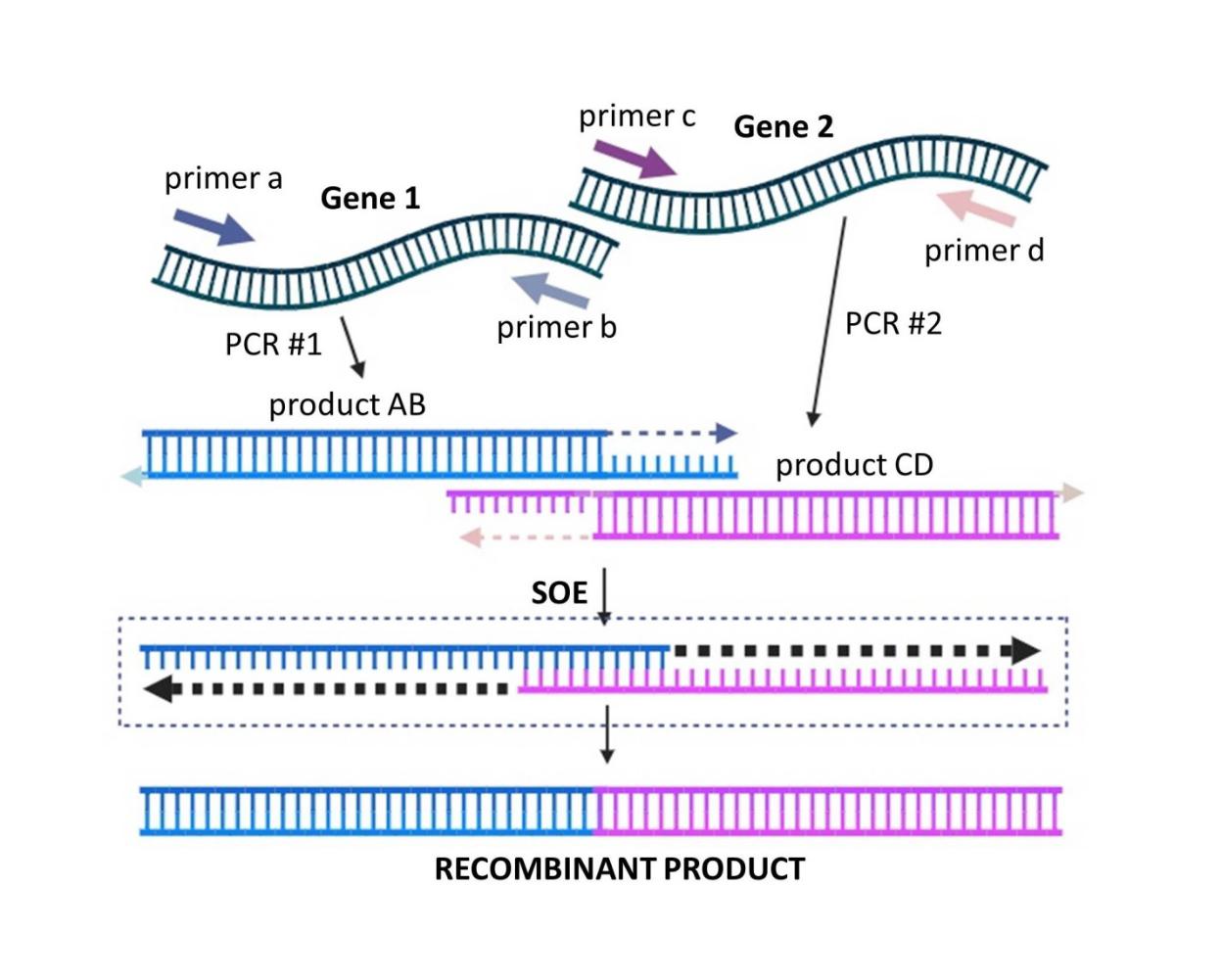

**SUPPLEMENTAL FIGURE 1 An illustration of the concept of SOE (gene Splicing by Overlap Extension).** PCR#1 amplifies Gene I fragment AB, and PCR #2 amplifies Gene II into the fragment CD. Primers **b** and **c** are the SOE primers, and they are used to modify the end of the two PCR products so that they have the same sequence. When these PCR products are mixed, denatured and reannealed under PCR conditions,

the top strand of AB and the bottom strand of CD overlap and act as primers on one another, leading to the formation of a recombinant product (as shown inside the broken rectangle). Inclusion of the outside primers **a**, **d** in the SOE reaction causes the recombinant product to be PCR amplified right after it is formed.

Apollo promoter variants were cloned by overlap PCR (CloneAmp proofreading enzyme) by using two rounds of PCR. In PCR round one (PCR1), promoter fragments were amplified from apomictic (ES517) or sexual (ES718) *Boechera* genomic DNA. For making constructs pC1 and pC2 (apomictic and sexual, respectively), the entire 2 kb promoter regions, including the native 5'UTRs, were amplified without alteration (Supplemental Table 2). Primers for these PCRs did not include overhangs, aside from the CACC tag used for TOPO directional cloning as shown in (Supplemental Table 5).

For pC3, pC4, and pC5 the 2 kb promoter region up to, but not including, the 20 bp Apo-insertion site contained within the 5'UTR, was spliced with an alternate version of the 5'UTR and/or Apo-insertion. To do this, the two spliced pieces of DNA were amplified separately (CloneAmp proofreading enzyme) with complementary overhangs incorporated into the internal primers (Supplemental Table 2). For pC3, primer overhangs were designed to “skip” amplification of the Apo-specific insertion, removing it from the final sequence. For pC4, primer overhangs incorporated the Apo-specific insertion as the overlapping region so that it would be incorporated into the final sequence. For pC5, primer overhangs were designed to incorporate a randomized insertion into the final promoter sequence.

The randomized insertion was created to check if any changes in the 20 bp Apo-insertion sequences order create a new GUS activity pattern. In order to make a randomized sequence the first step was to design the random 20 bp insertion using the following website (<https://www.bioinformatics.org/sms2/shuffle_dna.html>). Firstly, the 20 bp apomixis-specific polymorphism was entered as the raw sequence. Secondly, the randomized sequence was used to design the overlap primer the same was used for other constructs.

For pC6 and pC7, the 2 kb promoter region up to, but not including, the 5'UTR, was spliced with the alternate 5'UTR (Supplemental Table 3). As above, the two fragments were amplified independently with complementary overhangs incorporated into the primers. Beyond this splicing, no additional mutations or deletions were made.

In order to conduct the second round of PCR (PCR2), an approximately equimolar ratio of a pair of purified PCR products from the first round of PCR (PCR1) was created with the same forward and reverse primers used to create pC1 and pC2 to amplify the fully-spliced sequences.

**Construct pC3**

In order to make construct pC3, pAPOLLO AF -2 kb (internal) and pAPOLLOa -64R +oh3 (overhang) were used as primer pairs and ES517 was used as DNA sample to amplify the 2 kb upstream of the Apo promoter up to the 20 bp Apo-insertion/Sex-deletion site. Secondly, pAPOLLOa -44F +oh3 (overhang) and pAPOLLO 1R (internal) were used as primer pairs and ES 517 as DNA sample to amplify 44 bp of the 5'UTR of apo-allele promoter by removing the 20 bp Apo-insertion. Finally, to amplify the full spliced sequences an approximately equimolar ratio of a pair of purified PCR products from the first round one of PCR (PCR1) used as DNA samples for the second round of PCR (PCR2) with primer pairs of pAPOLLO AF -2 kb and pAPOLLO 1R.

**Construct pC4**

In order to make construct pC4 primer pairs of pAPOLLO SF-2 kb (internal) and pAPOLLOs - 58R +oh4 (overhang) and DNA of ES718 (Sex) were used to amplify the 2 kb upstream of Sex promoter by incorporating the 20 bp Apo-insertion. Secondly, the overhang primer pairs of pAPOLLOa -58F +oh4 (overhang) and pAPOLLO 1R (internal) and DNA of ES718 were used to amplify 52 bp of the 5’UTR of Sex promoter by incorporating the 20 bp Apo-insertion. Finally, to amplify the full spliced sequences an approximately equimolar ratio of a pair of purified PCR products from the first round one of PCR (PCR1) were used as DNA samples for the second round of PCR (PCR2) with primer pairs of pAPOLLO SF -2 kb and pAPOLLO 1R. In fact, the 20 bp Apo-insertion incorporated into the products of PCR1 (i.e., 4a & 4b) in Supplemental Table (3) were used as recombination sites for the PCR2 and eventually became part of the final product.

**Construct pC5**

In order to make construct pC5, primer pairs of pAPOLLO AF -2 kb (internal) and pAPOLLOa - 64R +oh5 (overhang) and DNA of ES517 (Apo) were used to amplify the 1,917 bp upstream of the Apo promoter by incorporating a random 20 bp insertion. Secondly, the primer pairs of pAPOLLOa -44F +oh5 (overhang) and pAPOLLO 1R (internal) and DNA of ES517 (Apo) were used to amplify 52 bp of the 5'UTR of Apo promoter by incorporating a random 20 bp insertion

(AAGTCTATAGCACGTGCATC). Finally, to amplify the full spliced sequences an approximately equimolar ratio of a pair of purified PCR products from the first round one of PCR (PCR1) were used as DNA samples for the second round of PCR (PCR2) with primer pairs of pAPOLLO AF -2 kb and pAPOLLO 1R. In fact, the 20 bp random insertion incorporated into the products of PCR1 (i.e., 5a & 5b) in Supplemental Table (3) were used as recombination sites for the PCR2 and eventually became part of the final product.

**Construct pC6**

In order to make construct pC6, pAPOLLO AF -2 kb (internal) and pAPOLLOa -71R +oh6 (overhang) were used as primer pairs and ES517 (Apo) as DNA sample to amplify the 1,870 bp upstream of the Apo promoter up to, but not including the 20 bp Apo-insertion/Sex-deletion. Secondly, primer pairs of pAPOLLOa -104F +oh6 (overhang) and pAPOLLO 1R (internal), and DNA sample of ES718 (Sex) were used to amplify 104 bp of the 5'UTR of Sex promoter. Finally, to amplify the full spliced sequences an approximately equimolar ratio of a pair of purified PCR products from the first round one of PCR (PCR1) were used as DNA samples for the second round of PCR (PCR2) with primer pairs of pAPOLLO AF -2 kb and pAPOLLO 1R.

**Construct pC7**

In order to make construct pC7, primer pairs of pAPOLLO SF -2 kb (internal) and pAPOLLOa - 104R +oh7 (overhang) and sample of ES718 (Sex) were used to amplify the 1,865 bp upstream of the Sex promoter up to, but not including the 5'UTR. Secondly, primer pairs of pAPOLLOa -71F +oh7 (overhang) and pAPOLLO 1R (internal), and DNA sample of ES518 (Apo) were used to amplify 71 bp of the 5'UTR of the Apo promoter. Finally, to amplify the full spliced sequences an approximately equimolar ratio of a pair of purified PCR products from the first round one of PCR (PCR1) used DNA samples for the second round of PCR (PCR2) with primer pairs of pAPOLLO SF -2 kb and pAPOLLO 1R.

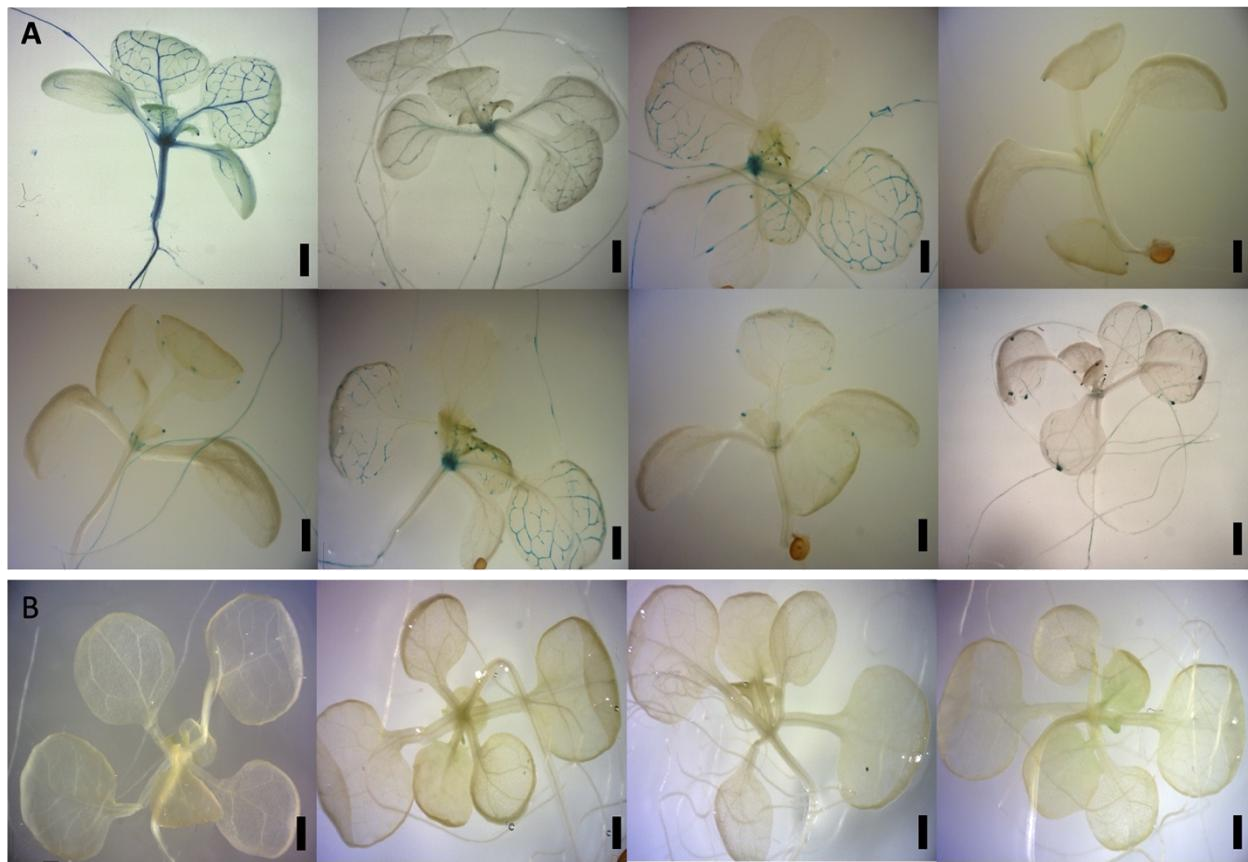

**SUPPLEMENTAL FIGURE 2.** Variation in expression patterns of *Boechera* native promoter GUS constructs in *Arabidopsis*. (A) GUS distribution of 2 kb native pApo promoter (pC1) in different replicates of two-week old plants. (B) No GUS activity observed in 2 kb native pSex promoter (pC2) in different replicates. (A,B) Scale bar=0.5 mm (Zeiss Lumar.V12 Stereoscope).

**(Grey value) GUS activity**

1

0.9

0.8

0.7

0.6

0.5

0.4

0.3

0.2

0.1

0

***AtpC3::GUS***

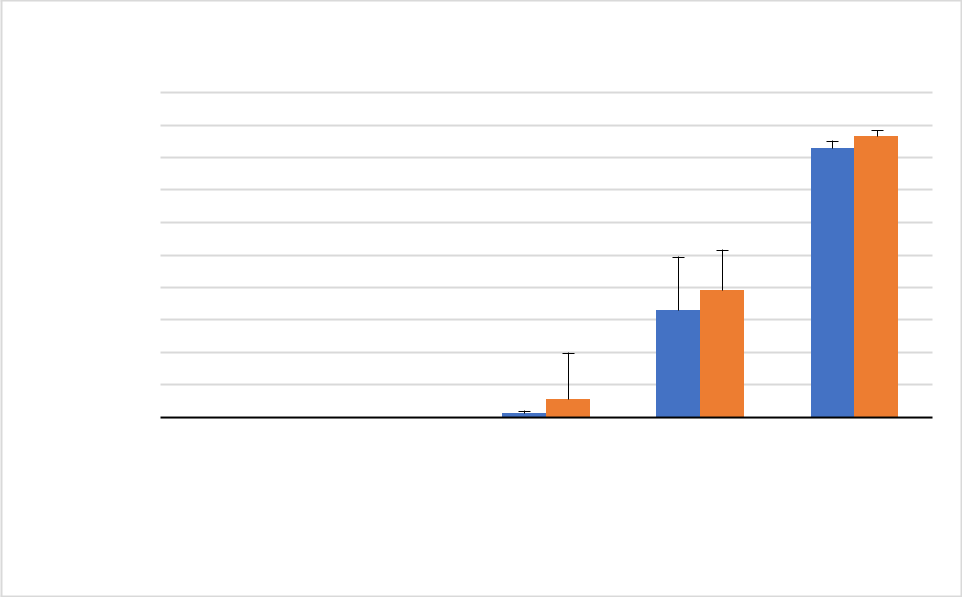

*** ^*^**

*** ^*^**

0.7mm	1.0mm	1.5mm	2.0mm	3.0mm

**Developmental Stages**

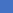
 Style
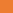
 Filament

**SUPPLEMENTAL FIGURE 3** Quantification of GUS histochemical staining in flowers of pC3: GUS transgenic *Arabidopsis*. Graph showing the mean GUS intensity (SD) measured in style and filament for five developmental stages within flowers. Asterisks indicate two final developmental stages are significantly different P≤0.05 from others (Tukey test; n=40 flower buds). The data were normalized using the Min-Max method. X-axis line represents different developmental stages of flower (based on the carpel height in mm). AtpC3: GUS means *Arabidopsis* transgenic for pC3: GUS.

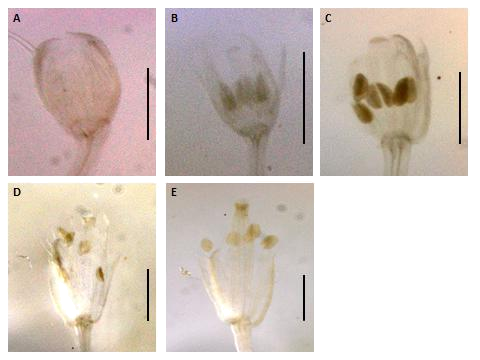

**SUPPLEMENTAL FIGURE 4** Expression patterns of spliced promoter construct (pC4: GUS) in *Arabidopsis* flowers. Construct pC4 promoters in different developmental stages of flowers. (A) 0.7 mm flower with no GUS activity, Scale bar=0.5 mm (B) 1.0 mm flower with no GUS activity. (C) 1.5 mm flower with no GUS activity. (D) 2.0 mm flower with no GUS activity. (E) 2.5 mm flower with no GUS activity. (B-E) Scale bar=1.0 mm (Zeiss Lumar.V12 Stereoscope).

**(Grey value) GUS activity**

1.1

1

0.9

0.8

0.7

0.6

0.5

0.4

0.3

0.2

0.1

0

***AtpC5::GUS***

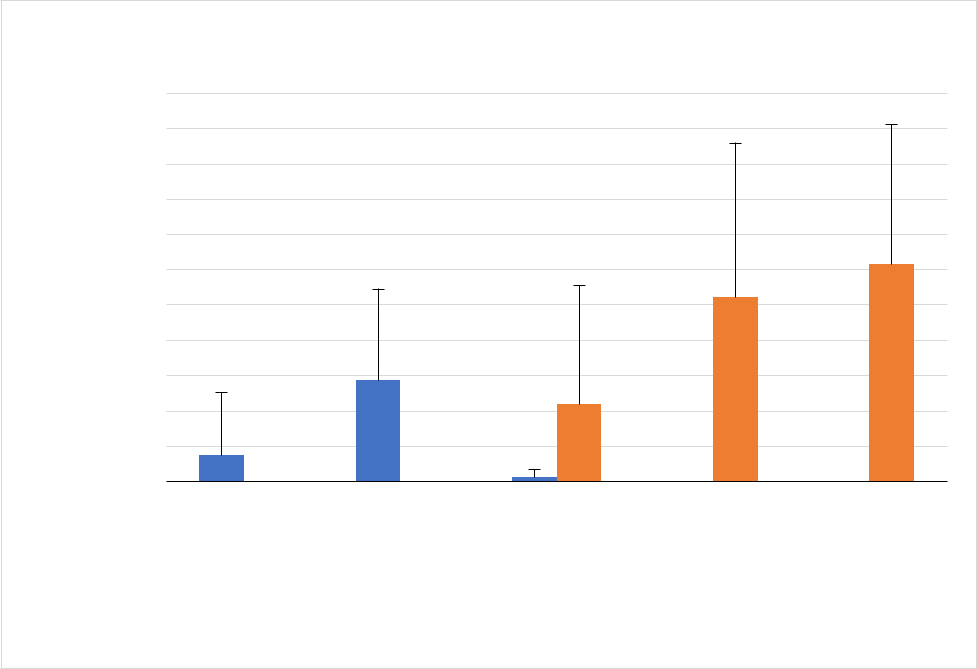

*****	**^*^**

0.7mm	1.0mm	1.5mm	2.0mm	3.0mm

**Developmental Stages**

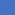
 Anther
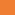
 Style

**SUPPLEMENTAL FIGURE 5** Quantification of GUS histochemical staining in flowers of pC5: GUS transgenic *Arabidopsis*. The image J quantification of GUS activity for construct (pC5) was carried out for 6 transgenic lines each of which had five different developmental stages (total of 30 flower buds). Graph showing the mean GUS intensity (SD) measured in different tissues (i.e., anther and style) and developmental stages within transgenic flowers. Asterisk indicates the last two developmental stages are significantly different P≤0.05 from others (Tukey test, n=90 flower buds). The data were normalized using the Min-Max method. X-axis line represents different developmental stages of flower (based on the carpel height in mm). AtpC5: GUS means *Arabidopsis* transgenic for pC5:GUS.

**(Grey value) GUS activity**

|  | ***AtpC6::GUS*** | | |  |  |  |  |  |  |  |  |  |  |  |  |  |
| --- | --- | --- | --- | --- | --- | --- | --- | --- | --- | --- | --- | --- | --- | --- | --- | --- |
| 1 |  |  |  | ***** | | | |  |  |  |  |  |  |  |  |  |
| 0.9 |  |  |  |  |  |  |  |  |  |  |  |  |  |  |  |  |
| 0.8 |  |  |  |  |  |  |  |  |  |  |  |  |  |  |  |  |
| 0.7 |  |  |  |  |  |  |  |  |  |  |  |  |  |  |  |  |
| 0.6 |  |  |  |  |  |  |  |  |  |  |  |  |  |  |  |  |
| 0.5 |  |  |  |  |  |  |  |  |  |  |  |  |  |  |  |  |
| 0.4 |  |  |  |  |  |  |  |  |  |  |  |  |  |  |  |  |
| 0.3 |  |  |  |  |  |  |  |  |  |  |  |  |  |  |  |  |
| 0.2 |  |  |  |  |  |  |  |  |  |  |  |  |  |  |  |  |
| 0.1 |  |  |  |  |  |  |  |  |  |  |  |  |  |  |  |  |
| 0 | 0.7mm | | | 1.0mm | | | | 1.5mm | | | | 2.0mm | | | | 3.0mm |

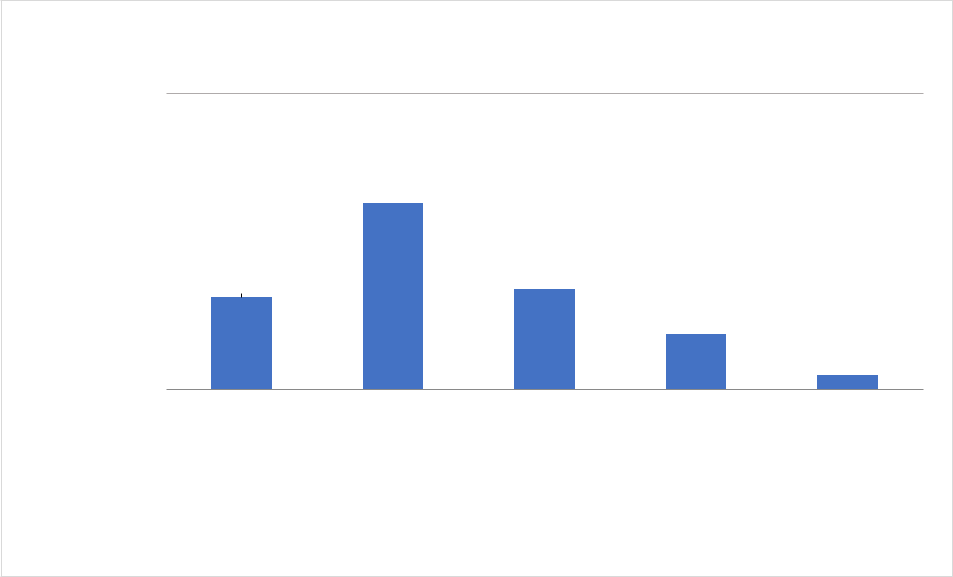

**Developmental Stages**

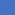
 Anther

**SUPPLEMENTAL FIGURE 6** Quantification of GUS histochemical staining in flowers of PC6: GUS transgenic *Arabidopsis*. Graph showing the mean GUS intensity (SD) measured in anther of different developmental stages. Asterisk indicates the second developmental stage is significantly different P≤0.05 from others (Tukey test; n=20 flower buds). The data were normalized using the Min-Max method. X-axis line represents different developmental stages of flower (based on the carpel height in mm). AtpC6: GUS means *Arabidopsis* transgenic for pC6: GUS.

**(Grey Value) GUS activity**

1

0.9

0.8

0.7

0.6

0.5

0.4

0.3

0.2

0.1

0

| ***AtpC7::GUS*** | | | | | | | | | |  |
| --- | --- | --- | --- | --- | --- | --- | --- | --- | --- | --- |
| *** * *** | | | | | | | | | | *** *** |

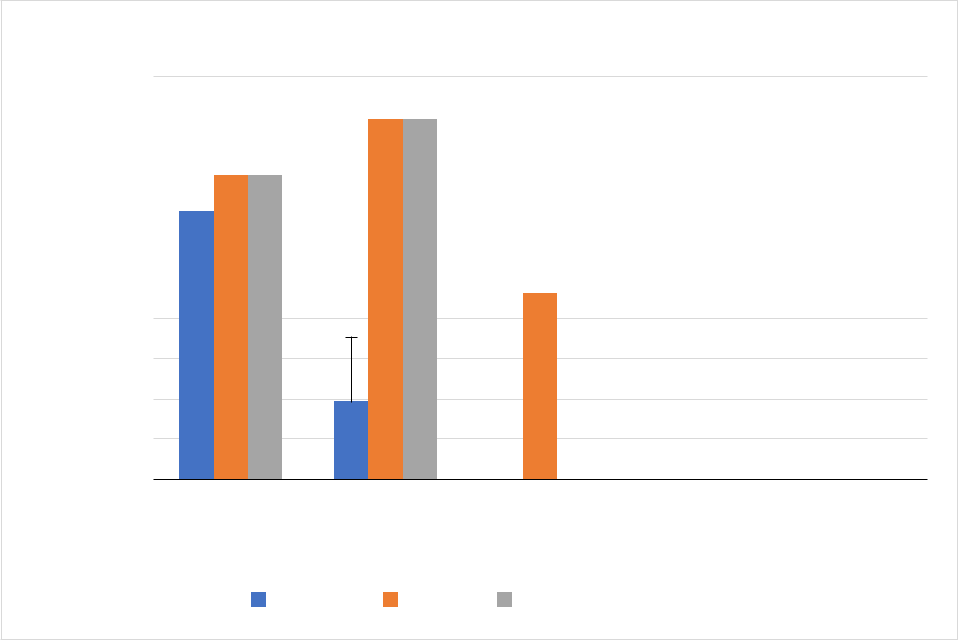

*****

| 0.7 mm | 1.0 mm | 1.5 mm | 2.0 mm | 3.0 mm |
| --- | --- | --- | --- | --- |
|  | **Developmental Stages** | | |  |
|  | AntherOvule | Transmitting Tract | |  |

**SUPPLEMENTAL FIGURE 7** Quantification of GUS histochemical staining in flowers of pC7: GUS transgenic *Arabidopsis*. Graph showing the mean GUS intensity (SD) measured in different tissues and developmental stages within transgenic flowers. Asterisks indicate the first two developmental stages are significantly different P≤0.05 from others (Tukey test; n=30 flower buds). X-axis line represents different developmental stages of flower (based on the carpel height in mm). The data were normalized using the Min-Max method. AtpC7: GUS means *Arabidopsis* transgenic for pC7:GUS.

**SUPPLEMENTAL TABLE 1** Native promoter (2 kb) and synthetic promoter constructs. Two native (pC1 and pC2) and five synthetic promoter constructs were created to test the importance of different apomictic-versus sex-specific polymorphism identified in *Boechera* (all have the same length of ~2 kb).

| **Construct name** | **Promoter type core** | **20 bp Apo-insertion** | **5' UTR** | **Transgenic lines** |
| --- | --- | --- | --- | --- |
| **pC1** | Apo promoter | **+** | Apo | *Arabidopsis/Boechera* |
| **pC2** | Sex promoter | **-** | Sex | *Arabidopsis/Boechera* |
| **pC3** | Apo promoter | **-** | Apo | *Arabidopsis* |
| **pC4** | Sex promoter | **+** | Sex | *Arabidopsis* |
| **pC5** | Apo promoter | Random insertion | Apo | *Arabidopsis* |
| **pC6** | Apo promoter | **-** | Sex | *Arabidopsis* |
| **pC7** | Sex promoter | **+** | Apo | *Arabidopsis* |

**SUPPLEMENTAL TABLE 2** Primer pairs and templates used for the first round of overlap PCR. Native constructs pC1 and pC2 were amplified using one set of primers, while two sets of internal and overhang primers were used for constructs pC3 to pC7. Constructs pC1 and pC2 only have internal primers.

| **PCR1** |  |  |  |  |  |  |
| --- | --- | --- | --- | --- | --- | --- |
| **Construct** | **F primer** | **R primer** | **Template** | **F primer** | **R primer** | **Template** |
| **pC1** | pAPOLLO AF -2 kb | pAPOLLO 1R | ES517 |  |  |  |
|  |  |  | (Apo) |  |  |  |
| **pC2** | pAPOLLO SF -2 kb | pAPOLLO 1R | ES718 |  |  |  |
|  |  |  | (Sex) |  |  |  |

|  | **Internal** | **Overhang** |  | **Internal** | **Overhang** |  |
| --- | --- | --- | --- | --- | --- | --- |
| **pC3** | pAPOLLO AF -2 kb | pAPOLLOa -64R +oh3 | ES517 | pAPOLLOa -44F +oh3 | pAPOLLO | ES517 |
|  |  |  | (Apo) |  | 1R | (Apo) |
| **pC4** | pAPOLLO SF -2 kb | pAPOLLOs -58R +oh4 | ES718 | pAPOLLOs -58F +oh4 | pAPOLLO | ES718 |
|  |  |  | (Sex) |  | 1R | (Sex) |
| **pC5** | pAPOLLO AF -2 kb | pAPOLLOa -64R +oh5 | ES517 | pAPOLLOa -44F +oh5 | pAPOLLO | ES517 |
|  |  |  | (Apo) |  | 1R | (Apo) |
| **pC6** | pAPOLLO AF -2 kb | pAPOLLOa -71R +oh6(s) | ES517 | pAPOLLOs -104F +oh6(a) | pAPOLLO | ES718 |
|  |  |  | (Apo) |  | 1R | (Sex) |
| **pC7** | pAPOLLO SF -2 kb | pAPOLLOs -104R +oh7(a) | ES718 | pAPOLLOa -71F +oh7(s) | pAPOLLO | ES517 |
|  |  |  | (Sex) |  | 1R | (Apo) |

**SUPPLEMENTAL TABLE 3** Primer pairs and templates used for (PCR2) of overlap PCR.

| **PCR2** |  |  |  |
| --- | --- | --- | --- |
|  | **F primer** | **R primer** | **Template** |
| **pC3** | pAPOLLO AF -2 kb | pAPOLLO 1R | PCR1 3a+3b |
| **pC4** | pAPOLLO SF -2 kb | pAPOLLO 1R | PCR1 4a+4b |
| **pC5** | pAPOLLO AF -2 kb | pAPOLLO 1R | PCR1 5a+5b |
| **pC6** | pAPOLLO AF -2 kb | pAPOLLO 1R | PCR1 6a+6b |
| **pC7** | pAPOLLO SF -2 kb | pAPOLLO 1R | PCR1 7a+7b |

**SUPPLEMENTAL TABLE 4** The Gene bank IDs for the promotor’s constructs.

**Construct name**	**Gene Bank ID**

pC1	ON668425

pC2	ON668426

pC3	ON668427

pC4	ON668428

pC5	ON668429

pC6	ON668430

pC7	ON668431

**SUPPLEMENTAL TABLE 5** Primer sequences used for overlap PCR. The CACC overhang of the first two primers and the overhangs of the other primers are highlighted in bold.

**Primer Name**

**Primer Sequence**

**pAPOLLO Apo F -2 kb**

**CACC**ACGAGGGATCCGGACAAAGATAC

**pAPOLLO Sex F -2 kb**

**CACC**CTCTGTTTCTTTCGTCCCGGTATTT

**pAPOLLO 1R**

TGTTAAGAACTGAGAGTGAAGGAG

**pAPOLLOa -44F +oh3**

**TTTTTTCCGTAAAAAGAGGAGG**CTTTAAAACCCACCAATTAGC

**pAPOLLOa -58F +oh4**

**TGGCCCGTGAAGTTTATTCC**ATCGATTGCTTTAAAACCCACC

**pAPOLLOa -44F +oh5**

**AAGTCTATAGCACGTGCATC**CTTTAAAACCCACCAATTAGC

**pAPOLLOa -64R +oh3**

**GCTAATTGGTGGGTTTTAAAG**CCTCCTCTTTTTACGGAAAAAA

**pAPOLLO -58R +oh4**

**GGAATAAACTTCACGGGCCA**CCTCCTCTTTTTACGGAAA

**pAPOLLOa -64R +oh5**

**GATGCACGTGCTATAGACTT**CCTCCTCTTTTTACGGAAAAAA

**pAPOLLOa -104F +oh6**

**TTTAGATTTTTTTCCGTAAAAA**TCGTACCGTTGCTTCTCTCAAG

**pAPOLLOa -71R +oh6**

**GAGAGAAGCAACGGTACGATTT**TTACGGAAAAAAATCTAAACTTG

**pAPOLLOa -71F +oh7**

**ATGACGCAAGATAAACCTCA**GAGGAGGTGGCCCGTGAAGTT

**pAPOLLOa -104R +oh7**

**AACTTCACGGGCCACCTCCTC**CTTGAGAGAAGCAACGGTACGA
